# Slow Oscillatory Transcranial Direct Current Stimulation during a Restricted Sleep Opportunity Enhances Cognitive Performance during Subsequent Wakefulness

**DOI:** 10.64898/2026.07.03.736438

**Authors:** John D. Hughes, Tracy Jill Doty, Thomas J. Balkin

## Abstract

The slow oscillation (SO) of non-rapid eye movement (NREM) sleep has been implicated in the restorative properties of sleep. Slow oscillatory transcranial direct current stimulation (SO-tDCS), involving a positive oscillatory current applied to the scalp at a peak frequency of 0.75 Hz, has been used to enhance SO power during NREM sleep. We examined whether enhancing SO power with SO-tDCS during a restricted nighttime sleep opportunity would accelerate the restorative properties of sleep during an otherwise insufficient sleep period and help sustain performance during subsequent extended wakefulness.

A total of twenty-six healthy young adults (ages 18-39, n=16 females) completed a 15-day study. After 7 baseline nights at home and 3 baseline nights in the laboratory, participants entered the laboratory for 5 consecutive days including a baseline day, a 2-hour nighttime sleep period with participants randomized to the SO-tDCS (n=11) or SHAM (n=15) condition, 46 hours of sleep deprivation, and two recovery nights. In the SO-tDCS condition, stimulation was administered for one hour starting exactly 60 minutes after sleep onset, with intervals of five minutes of continuous stimulation followed by one minute of no stimulation.

Polysomnographic recordings were conducted during each sleep period. Performance was assessed using the Psychomotor Vigilance Test (PVT) approximately every 75 minutes across baseline, sleep deprivation, and recovery. Prior to the two-hour sleep opportunity, a Paired Words Associate Task was administered. Participants listened to 54-word pairs and were asked to recall 46 of the word pairs, with up to three attempts to successfully recall at least 60% of word pairs (T0). Recall was also assessed 20– (T20) and 120-minutes (T120) after awakening from the two-hour sleep period. Data were analyzed using mixed-effects ANOVA.

PVT performance (defined as mean response time and number of response times greater than 1,000 ms) significantly declined across sleep deprivation with performance degradations peaking in the early morning hours. Participants in the STIM condition demonstrated significantly better performance during sleep deprivation relative to the SHAM condition. On the PWAT, participants in the SHAM condition recalled fewer word-pairs upon awakening relative to T0. In sharp contrast, performance of participants in the SO-tDCS condition did not deteriorate at T20 and was actually improved at T120 relative to T0.

We conclude that SO-tDCS can robustly accelerate sleep’s restorative processes and can additionally enhance sleep related memory consolidation when sleep opportunity is restricted.

## Introduction

Non-rapid eye movement (NREM) sleep electroencephalogram (EEG) is characterized by the presence of high amplitude slow-wave activity. Such activity was first identified in 1929 by EEG inventor Hans Berger and further characterized in the 1930s by sleep physiologist Alfred Loomis. However, for more than 60 years slow-wave activity was viewed as simply a passive idling rhythm which emerges when excitation of the neocortex via ascending reticular formation activation is reduced and metabolically more demanding fast, low amplitude EEG activity consistent with conscious cognition is turned off, with the cortex entering a state of “rest” and restoration (Steriade, 1970). Even when a restorative function of deep slow-wave *sleep* was being proposed in the 1970s and 1980s (Adam 1980, Spiegel 1986), an active mechanistic role of the slow waves themselves was not part of these hypotheses. However, in the last two decades, as the physiology of sleep slow waves and related EEG rhythms have been elucidated, important sleep related functions have been increasingly attributed to the actual slow waves, rather than to an absence of faster waves. Slow waves with a frequency range of 0.3 to 4 Hz, traditionally referred to as delta frequency activity, consist of different components with unique physiological underpinnings. In the early 1990s, Steriade and colleagues first characterized the physiology of activity in the slowest component of delta activity, the 0.3 to 1.0 Hz range, which they referred to as the “< 1 Hz slow oscillation”, as consisting of a recurrent alternation between two stable states of neuronal membrane potential, a more depolarized one (the UP state) and a more hyperpolarized one (the DOWN state), and as being distinct from the rest of higher frequency delta activity (Steriade et al, 1993a). This activity constitutes the majority of the EEG power in the slow-wave range, is generated independently in the cortex, though modulated by a more rhythmic SO generated independently in the thalamus, and has been the focus of slow-wave research in recent years. The remainder of EEG delta activity, with a frequency range of approximately one to four Hz, has multiple proposed physiological origins in both the cortex and thalamus, and, with a faster frequency, can be superimposed on either the UP or DOWN state of the slow oscillation (Steriade, 1993b; Hughes et al, 2002) or occur independently.

The slow oscillation, including its specific phasic oscillatory components, has been hypothesized to play an active mechanistic role in two important but separate proposed functions of non-rapid eye movement (NREM) sleep: sleep-related declarative memory consolidation and *sleep restorative function*. With regard to memory consolidation, the transition from the DOWN to the UP state of the slow oscillation, transmitted throughout cortex and from cortex to subcortical structures, is thought to trigger and synchronize thalamocortical sleep spindles and hippocampal sharp-wave ripple activity that support the transfer of declarative memories from their initial encoding site in the hippocampus to more permanent representation in the neocortex, integrated into existing memory networks (Buzsaki, 1989). This transfer of encoded memory representations from a distributed representation based in the hippocampus to a distributed cortical representation free of a hippocampal component is referred to as *systems consolidation*; the slow oscillation is the foundational physiological basis for this process (Lutz et al, 2015).

Additionally, the slow oscillation is believed to actively mediate sleep’s restorative function. By restorative function, we refer to the process or processes by which sleep reverses the deleterious effect of sustained wakefulness on performance, and the associated accrual of homeostatic sleep drive. A leading theory of the restorative function of sleep, the synaptic homeostasis hypothesis (SHY) of Tononi and Cirelli, purports that that the recurrent alteration between UP and DOWN states of the slow oscillation actively downscales synapses throughout the cortex, thereby reducing overall cortical synaptic strength that increases gradually during sustained wakefulness, due to continuous information processing with Hebbian plasticity mechanisms in effect (Tononi and Cirelli, 2006; 2014, 2020). The increase in global synaptic strength renders cortical processing inefficient due to an imbalance between neuronal excitation and inhibition, leading to reduced attentional and related cognitive functions, saturation of learning ability and presumably sleepiness. Another prominent theory (Vyazovskiy and Harris, 2013) proposes that the DOWN state of the SO contributes to sleep’s restorative properties by providing a hyperpolarized neuronal state devoid of dendritic synaptic activity and therefore somatic electrical silence, which allows neuronal cellular repair of damage to DNA and organelles incurred during the metabolically demanding activity of generation of repetitive action potentials and which would lead to cell apoptosis if such damage continued to accumulate unabated. The electrical silence of the DOWN state is felt to be essential for this repair to take place. Finally, a third hypothesis suggests that the SO (and more specifically DOWN states) enhances the elimination of toxic byproducts of cellular metabolic activity from the brain parenchyma, via the brain’s “glymphatic” system (Xie et al, 2003). Based on this theory, at least some component of the accumulation of homeostatic drive is due to an increasing concentration of these products that disrupt normal neuronal function. These three theories of SO mechanisms of restorative function are in no way mutually exclusive and are likely complementary.

The proposed role of sleep slow waves in these important sleep restorative functions has sparked an interest in slow oscillation enhancement strategies both due to the growing epidemic in insufficient sleep in healthy individuals (Chattu et al, 2018) as well as the reduction in slow wave activity in several common neuropsychiatric disorders, including Alzheimer’s Disease (Lee et al, 2020). Pharmacological slow wave enhancement is of limited practical value due primarily to medication side effects and drug related tachyphylaxis. Non-invasive brain stimulation methodologies hold promise for slow-wave enhancement. In a seminal study, Marshall et al. (2006) demonstrated that 25 minutes of a slow oscillatory form of continuous transcranial direct current stimulation (SO-tDCS) with a frequency at approximately the peak of endogenous SO activity (0.75 Hz), during early nocturnal sleep, modestly improved subsequent retention of word pairs learned prior to sleep, consistent with the proposed role of the slow oscillation in sleep-related memory consolidation. Generation of individual slow waves with acoustic stimulation of discrete tones has also been demonstrated to enhance slow wave power to some degree and improve some aspects of sleep-related declarative memory consolidation with variable success (Ngo et al, 2013). However, acoustic stimulation is currently limited to stimulation in deep NREM sleep, already high in SO power, thus limiting the ability to enhance such power. The efficacy of SO-tDCS in enhancing the restorative function of sleep has not heretofore been assessed.

In this study, we sought to determine the efficacy of SO-tDCS induced enhancement of slow oscillation activity during a portion of a restricted period of sleep, prior to an extended period of sleep deprivation, to enhance the restorative properties of that sleep period via exogenous SO enhancement and thus render those individuals stimulated relatively resilient to the deleterious effects of sleep deprivation on alertness. Specifically, we assessed whether or not fifty minutes of SO-tDCS during a restricted two hour period of nocturnal sleep prior to a 46 hour period of sleep deprivation would render participants resilient to the effects of sleep deprivation on performance compared to a participants receiving sham stimulation. The very restricted period of sleep prior to a period of total sleep deprivation would ensure that study participants would be especially vulnerable to the deleterious effects of sustained wakefulness on cognitive performance; the degree to which those receiving SO-tDCS may be resilient to these detrimental effects would be evidence of an enhanced level of restorative physiology provided by the SO-tDCS.

Specifically, to assess the efficacy of SO-tDCS in enhancing the restorative properties of sleep, we subjected healthy participants to a restricted two hour period of nighttime sleep after a normal sixteen hour period of daytime sustained wakefulness with or without a 50 minute period of SO-tDCS during the second hour of sleep. We know that a two hour period of sleep is drastically insufficient to satisfy the accumulated homeostatic sleep drive from a sustained sixteen hour period of wakefulness. In fact even as little as one hour less than the optimal seven to eight hours of sleep will result in significant decrements in performance during subsequent wakefulness in the majority of individuals. Participants were subjected to a 46 hour period of total sleep deprivation (SD) immediately after the two hour period of restricted sleep. They were therefore entering SD with an already significant sleep debt and were therefore exceptionally vulnerable to the decrementing effects of SD on performance. Consequently, to the degree that the fifty minute period of SO-tDCS during the two hour period of restricted sleep would enhance the restorative properties of that sleep period, the group that received stimulation would exhibit preserved performance compared to the sham group. That is, the better the stimulation group would perform compared to the sham group during SD, the more effective the stimulation would be in enhancing the sleep related reduction in homeostatic sleep drive.

## Methods

### Participants

Healthy civilian and active-duty military men and women aged 18 to 39 years (inclusive) were recruited via an advertisement distributed to local universities and military installations. After providing informed consent and completing eligibility questionnaires, participants underwent general physical and neurological examinations as well as an evaluation of blood and urine samples to determine general health, including pregnancy and drug use. In order to reduce intersubject variability related to sleep habits, individuals were excluded from the study if they reported the following: (1) habitual nightly sleep amounts outside of 6–9 hours, (2) nighttime lights-out times earlier than 09:00 pm or later than midnight on average during weeknights, (3) morning wake-up times later than 09:00 am on average during weekdays, (4) habitual napping (>1 time a week in conjunction with normal sleep habits), an estimated sleep onset time of greater than 30 minutes and/or (5) a score lower than 31 and higher than 69 on the Morningness-Eveningness Questionnaire. To reduce variability in stimulant use and metabolism, individuals were excluded from the study if they reported caffeine use in excess of 400 mg per day on average and/or regular nicotine use (defined as more than one cigarette or equivalent per week) or if they had a body mass index of 30 or greater (classified as “obese”). Additionally, participants were excluded for use of tobacco/nicotine products in the last year or for more than moderate alcohol intake. Participants were also excluded if they had previously been diagnosed with any cardiovascular, pulmonary, neurological, kidney, liver, or psychiatric diseases and/or if they scored higher than 14 on the Beck Depression Inventory or higher than 41 on the Spielberger Trait Anxiety Inventory. Only participant who learned English as their first language were included in the study. With the exception of oral birth control pills, no participants were taking any prescription or over the counter medications. After eligibility was determined, participants were randomly assigned to one of two groups: stimulation (STIM group – 12 participants) or sham stimulation (SHAM – 15 participants). Mean ages per group: STIM: 25.1, SHAM 26.5. Due to the rigorous nature of our general and sleep health screen, 454 participants who responded to flyers recruiting “completely healthy” subjects were screened in order to achieve our final enrollment.

## Testing Facilities

During sleep satiation, baseline (BL), restriction, and recovery periods, each participant slept in their own sound-attenuated 8’ × 10’ room. Testing was completed in a designated testing area. When not engaged in testing or sleep, participants remained in a common living area to read, eat, play games, or watch television and movies. Lighting was set at <50 lux during waking hours so as to minimize the impact of light on circadian rhythms.

## Study Design

Participants underwent a 15 day protocol which included a 7 day actigraphy period, a 3 night sleep satiation period, a baseline day followed by a 2 hour period of sleep restriction, during which participants either received SO-tDCS stimulation or sham stimulation (a between subjects design). 2 days (a total of 46 hours) of sleep deprivation, and finally, two eight hour recovery nights followed by two recovery days. Due to the intense time demands of each stim or sham study component, which included a period of 5 continuous days and nights in our sleep laboratory without contact with the outside world, a within subjects cross-over design (initially intended) was not practical for the vast majority of potential participants who could not commit to two such components within a several month period of time.

**1.** **At-home sleep monitoring phase (7 nights):** Participants were instructed to maintain their chronic sleep schedule reported at screening. The participants’ sleep-wake activity was be assessed using actigraphy to ensure adherence to this requirement. Participants were required to refrain from taking daytime naps or study-prohibited substances (caffeine and alcohol) during this period.
**2.** **In-laboratory sleep satiation phase (3 nights**): Participants arrived at the laboratory nightly at approximately 1800. Participants had 10 hours in bed with lights out (2100 to 0700) and left the lab after 0700 after the first 2 nights (the day after the third sleep satiation night was the first night of the in lab phase-baseline day). Participants were instructed not to nap during the day and were monitored throughout this phase with actigraphy. During each night, participants underwent polysomnographic monitoring. The purpose of the sleep satiation phase was to attempt to align the circadian rhythms of the subjects as much as possible, and also to reduce or eliminate any chronic sleep debt that participants may have accumulated prior to study participation that may have otherwise introduced a confounding variable to the study data.
**3.** **In-laboratory sleep restriction/sleep deprivation phase (5 days/4 nights):** Participants remained in the lab the day after their last sleep satiation phase night, prior to their night of sleep restriction with stimulation intervention (STIM or SHAM). Baseline daytime performance assessment with the PVT took place every 75 minutes. Prior to bedtime on the sleep restriction night (following the baseline day), 4 transcranial stimulation electrodes (two on each hemisphere) were applied to the scalps of both groups (STIM and SHAM). Participants had a two hour sleep opportunity, with lights out at approximately 11:00. Sleep was monitored with EEG and after approximately one hour from the onset of the first period of stage N2 sleep that was sustained for at least 2 minutes, participants in the STIM group received a one hour period of slow oscillation transcranial direct current stimulation (SO-tDCS), for a total of approximately two hours of sleep. The participants in SHAM group received 60 minutes of stimulation at 0 current in place of the SO-tDCS. Participants in both groups were be awakened 2 hours after the onset of the initial period of stage N2 sleep, at the end of the 60 minute period of stimulation or sham stimulation.. Participants who experienced less than a total of approximately 30 minutes of sleep during the first of the 2 hour sleep opportunity, or experienced wakefulness during more than three of the one minute interstimulus intervals (including the one minute period after the last five minute stimulation block during the second of the two hour period during which stimulation or sham stimulation took place), were to be withdrawn from further involvement in the study. Additionally, those participants who had not fallen asleep within approximately 75 minutes of lights off (bedtime) were also excluded from further involvement in the study and their data was not analyzed. If a subject was awake and perceived a stimulation induced sensation on the scalp (tingling, paresthesias), he/she was allowed to remain in the study as long as he/she could fall back to sleep. However, he/she must be asleep for at least half of the stimulation period in order to continue with study participation. The two hour period of sleep restriction was followed by a 46 hour period of total sleep deprivation (TSD), during which performance was periodically assessed with the Psychomotor Vigilance Test (PVT). During the recovery sleep phase immediately following the period of TSD, subjects obtained two nights of recovery sleep consisting of 8 hours’ time in bed (from 11:00pm to 7:00 am). Performance assessments also occurred periodically on the day following each of the recovery nights of sleep. During the recovery nights, sleep was objectively monitored using actigraphy and polysomnography. Subjects were dismissed from the study at approximately 5:00 pm on the day following the second recovery night of sleep.

Sleep onset (time to start of first two minute period of continuous N2 sleep) varied from 14 minutes to 53 minutes among all study participants on the sleep restriction/stimulation intervention night. All participants were awakened 2 hours after the start of the first two minute period of continuous N2 sleep in order to begin the sleep deprivation phase of the study. Because sleep onset varied among participants, the following period of sleep deprivation also varied somewhat since recovery sleep started at the same time for all participants. Participants received between approximately 45 and 46 hours of SD. The average sleep onset for the STIM group was 27 minutes and for the SHAM group was 29 minutes. Therefore, the STIM group participants received on average 45 hours and 33 minutes of SD and the SHAM group received an average of 45 hours and 31 minutes of SD.

### Stimulation

The stimulator utilized in this study was the neuroConn DC stimulator plus MC. SO-tDCS consisted of a trapezoidal shaped waveform with 0.33 second periods of stimulation at 0 amperes and 260 microcamperes, with 0.33 second transitions between these high and low current plateaus, resulting in a 0.75 Hz oscillation between 0 and 260 microamperes, based on the stimulation paradigm in Marshall et al (2006). The stimulation sequence consisted of ten 5 minute periods of continuous stimulation and nine one minute interstimulus intervals between these five minute stimulation periods, for a total of fifty minutes of SO-tDCS or a sixty minute total stimulation period. One anode/cathode pair was placed on each hemisphere, with anodes adjacent to the recording electrodes atF3 and F4 and cathodes at M1 and M2 of the International 10:20 system. Preliminary work with this stimulation sequence in individuals while and asleep awake revealed that this weak current electrical stimulation sequence could rarely be minimally perceived while awake and could never be perceived while asleep (and did not arouse sleeping individuals). Debriefing after stimulation revealed that no participants had a sense as to whether or not they had received stimulation or sham stimulation. The beginning of the two hour sleep period (sleep onset) was set at the end of by the first two minute period of continuous N2 sleep. Stimulation or sham stimulation was initiated exactly 60 minutes after the sleep onset, at the beginning of the second hour of sleep, regardless of the amount or sleep obtained in that first 60 minutes (although excessive wakefulness after sleep onset would lead to study withdrawal as described), unless a participant was awake at that time. Sleep (NREM or REM) had to be present in order to initiate stimulation. If wakefulness was present, stimulation initiation was delayed until the first epoch of sleep of any type. Stimulation did not exceed the two hour period from sleep onset, regardless of when it was initiated with respect to the 60 minute point after sleep onset. However, continuous wakefulness of at least 15 minutes at the onset of the intended stimulation period resulted in study withdrawal. Additionally, subsequent five minute stimulation blocks were initiated regardless of brain state (wakefulness or sleep), although per the protocol, participants were to be withdrawn form study participation if more than three periods of one minute interstimulus intervals consisted of predominantly wakefulness. Therefore, per the protocol guidelines, participants were to receive between seven and ten five minute stimulation blocks depending on brain state during the beginning of the second hour of sleep. In reality, all 27 participants who completed the study received the entire ten intended blocks of stimulation or sham stimulation. However, not all participants received ten full stimulation blocks during sleep as determined by the presence of wakefulness at the beginning of some interstimulus intervals. Given the presence of high amplitude stimulation artifact, there is no way to be sure exactly what percentage of time during stimulation was spent in N2 or N3 sleep. The sham group spent on average of x minutes in N2 or N3 sleep during their analogous 5 minute stimulation blocks.

### Measurements

A 10-minute Psychomotor Vigilance Test (PVT) was administered approximately every 75 minutes while awake during baseline, sleep deprivation, and recovery (Dinges & Powell, 1985). Mean response time and number of major lapses (defined as reaction times exceeding 1,000 milliseconds) were calculated for each task administration. Average mean response time and number of major lapses were calculated for each study day. Data were analyzed using mixed effects ANOVA with a main effect of study day (five levels: baseline, sleep deprivation day 1, sleep deprivation day 2, recovery day 1, and recovery day 2) and condition (two levels: SHAM and SO-tDCS), their two-way interaction, and a random intercept over subject. Post-hoc comparisons were conducted to assess changes across study days and conditions. Analyses were performed using SPSS (Version 28; SPSS Inc., Chicago, IL). Variance is expressed as standard errors.

A Paired Word Associate Task (PWAT) was administered. Beginning one hour prior to the two-hour sleep period, participants listened to 54 semantically related or unrelated nouns at a rate of one word pair per 5 seconds. Participants were then asked to recall 46 of the word pairs (excluding the first and last four pairs to limit primacy and recency effects) and had up to three attempts to correctly recall at least 60% of the word pairs (T0). Recall was again tested at 20 (T20) and 120 minutes (T120) after awakening from the two-hour sleep period. The number of word pairs correctly recalled at T20 and T120 relative to T0 were analyzed with a mixed-effects ANOVA that included a main effect of group (two levels: SHAM and SO-tDCS) and timepoint (two levels: T20 and T120), their two-way interaction, and a random intercept over subjects. In this analysis, negative values indicate that subjects recalled fewer pairs post-awakening relative to initial learning recall.

## Results

A total of twenty-six participants completed the study (age: 25.6 ± 12.3 years, n=16 females), including 11 participants in the SO-tDCS condition (age: 27.3 ± 12.8 years, n=7 females) and 15 participants in the SHAM condition (age: 24.6 ± 12.3 years, n=4 females).

Using the Psychomotor Vigilance Test (PVT), there was a significant main effect of study day (number of lapses: F_4,1621_=77.16, p<0.0001; mean response time [RT]: F_4,1621_=18.67, p<0.0001), main effect of condition (number of lapses: F_4,1621_=21.61, p<0.0001; mean RT: F_4,1621_=18.67, p<0.0001), and interaction between day and condition (number of lapses: F_4,1621_=4.30, p=0.002; mean RT: F_4,1621_=6.84, p<0.0001) on performance. Post-hoc comparisons demonstrate that participants in the SO-tDCS condition exhibited significantly better performance during sleep deprivation than those in the SHAM condition. Using number of lapses, this effect persisted into the first recovery day, with participants receiving SO-tDCS exhibiting better performance than those in the SHAM condition with one recovery night. See Figures 1-2 & Table 1.

**Figure 1:**
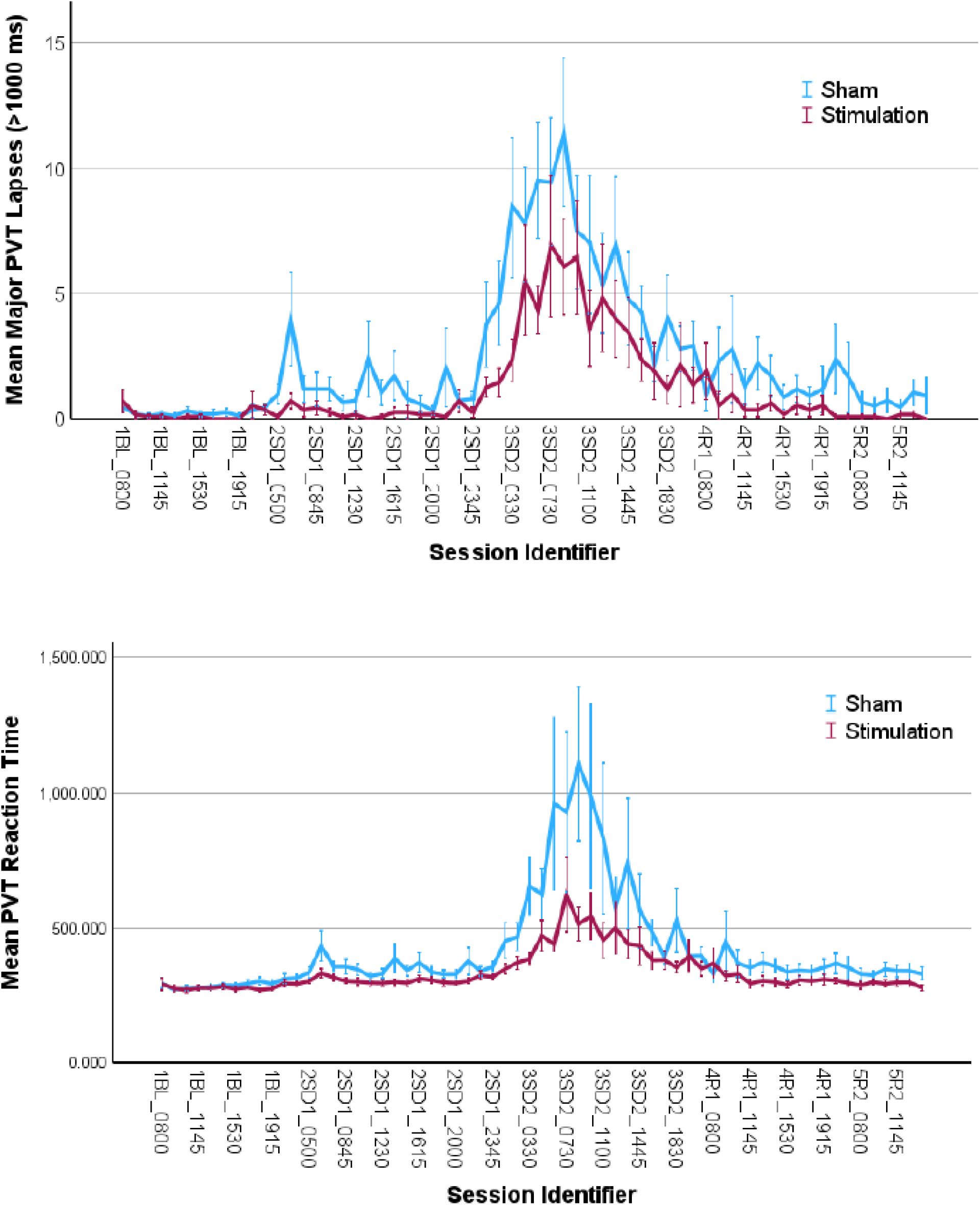
Psychomotor Vigilance Test (PVT) number of major lapses and mean RT (± standard error) for the SO-tDCS condition (“stimulation”, **red**) and SHAM condition (“Sham”, **blue**) across baseline (1BL), two days of sleep deprivation (2SD, 3SD), and two days of recovery (4R, 5R) at different times of day.

**Figure 2:**
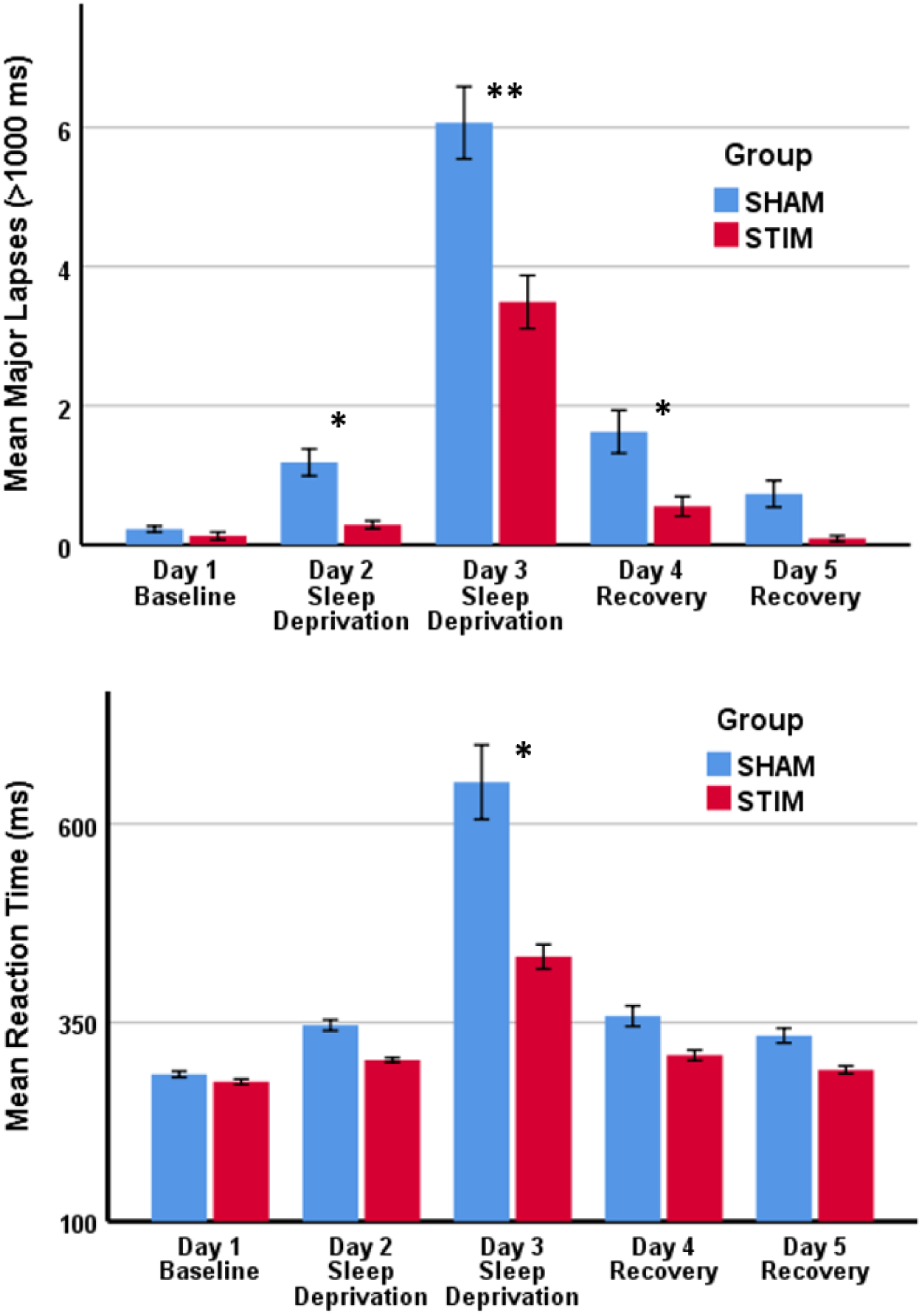
Average number of major lapses (top) and mean response time (bottom) for the SHAM (blue) and STIM/SO-tDCS (red) conditions across baseline, sleep deprivation, and subsequent recovery.

**Table 1.** Psychomotor Vigilance Test Performance.

| **Lapses (# RT > 1,000 ms)** |  |  |  |  |
| --- | --- | --- | --- | --- |
|  | **SHAM** |  | **SO-tDCS** |  |
| Study Day | Mean | SE | Mean | SE |
| 1BL | 0.23 | 0.34 | 0.13 | 0.40 |
| 2SD 1 | 1.19 | 0.26 | 0.29 | 0.30 |
| 3SD 2 | 6.07 | 0.26 | 3.49 | 0.31 |
| 4RC 1 | 1.63 | 0.32 | 0.55 | 0.36 |
| 5RC 2 | 0.73 | 0.44 | 0.09 | 0.52 |

| **Mean RT (ms)** |  |  |  |  |
| --- | --- | --- | --- | --- |
|  | **SHAM** |  | **SO-tDCS** |  |
| Study Day | Mean | SE | Mean | SE |
| 1BL | 285.03 | 25.40 | 275.61 | 29.66 |
| 2SD 1 | 346.85 | 18.93 | 303.09 | 22.11 |
| 3SD 2 | 652.51 | 19.59 | 432.96 | 22.75 |
| 4RC 1 | 358.28 | 23.45 | 308.96 | 27.07 |
| 5RC 2 | 333.74 | 32.79 | 290.65 | 38.29 |

\*Abbreviations: Baseline (BL), Sleep Deprivation (SD), Recovery (RC), Response Time (RT), Milliseconds (ms), Sham Stimulation (SHAM), Slow-Oscillatory Transcranial Direct Current Stimulation (SO-tDCS), Standard Error (SE)

On the Paired Words Associate Task (PWAT), participants correctly recalled 35.0 ± 9.2 words (mean ± standard deviation) following 2.1 ± 0.5 attempts during initial learning recall (T20). Following three attempts, one subject in the SHAM condition and one subject in the STIM condition had not correctly recalled at least 60% of the word-pairs. There was a significant main effect of group (F=6.20, p=0.02) and main effect timepoint (F=4.67, p=0.04) but not their two-way interaction (F=0.16, p=0.69). Relative to T0, subjects in the SHAM condition recalled significantly fewer word-pairs post-awakening than those in the SO-tDCS condition (SHAM: –0.8 ± 1.6 words; SO-tDCS: 0.3 ± 1.3 words), and significantly fewer word-pairs at T20 than T120 (T20: –0.6 ± 1.5 words; T120: –0.1 ± 1.6 words). On average, subjects in the SHAM condition recalled fewer word-pairs upon awakening relative to T0. In sharp contrast, performance of the SO-tDCS group did not deteriorate at T20 and was actually improved at T120. See Figure 2.

**Figure 2:**
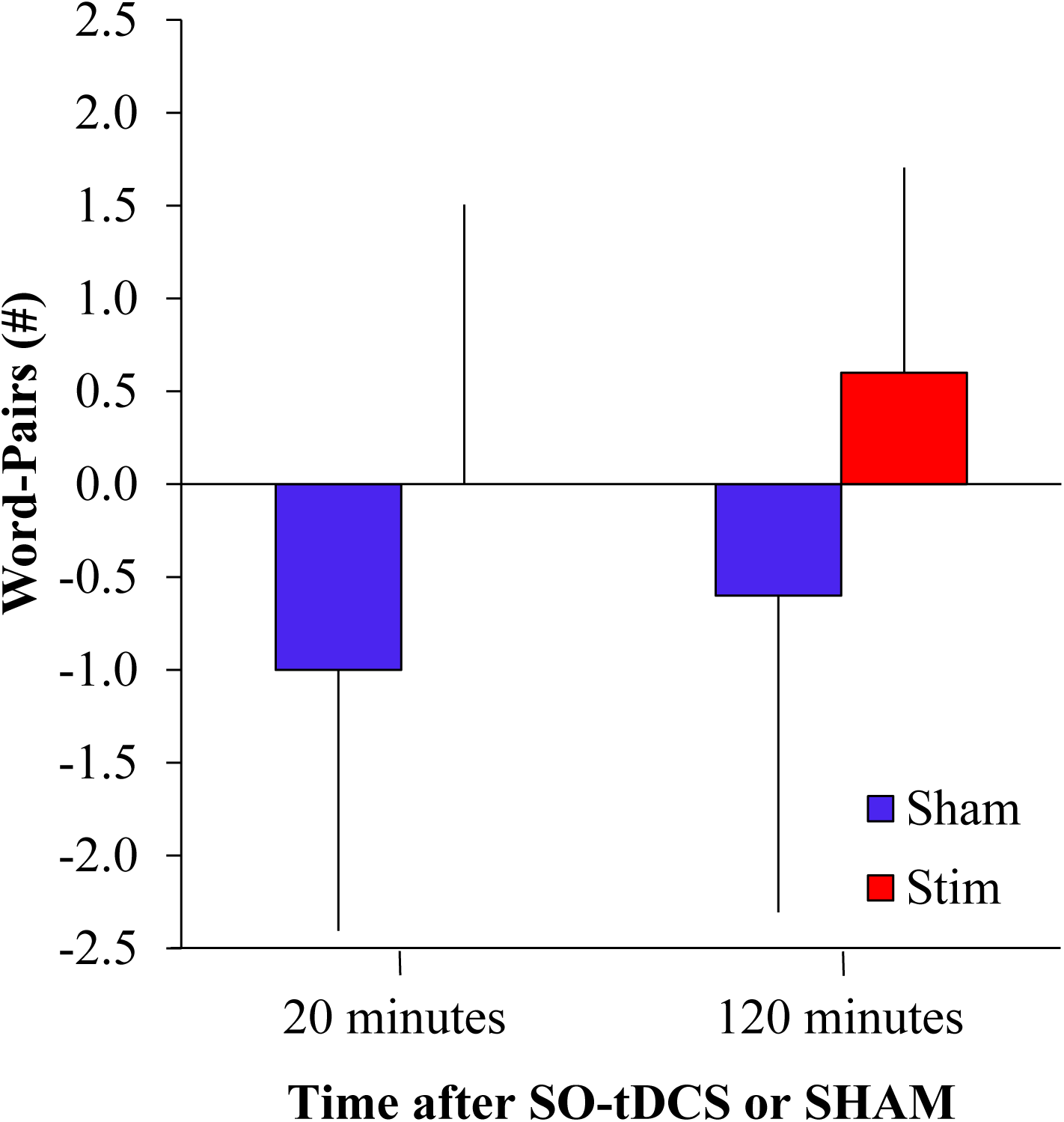
On the Paired Words Associate Task (PWAT), average number of word-pairs (± standard deviation) at 20– (T20) and 120– (T120) minute post-awakening relative to pre-sleep learning recall (T0) for SHAM (**blue**) and SO-tDCS (**red**) conditions. Negative values indicate that subjects recalled fewer word pairs post-awakening (T20 or T120) relative to the initial learning recall (T0).

## Discussion

In this study, we have demonstrated a robust effect of a single 50 minutes of SO-tDCS versus sham stimulation, during a restricted period of sleep, on performance during a subsequent period of sleep deprivation and a recovery period, as assessed with the PVT, a tool known to be exquisitely sensitive to the deleterious effects of sleep loss on performance. Using both PVT reaction time and number of major lapses, the STIM group outperformed the SHAM group at all time-points after the stimulation period including the two days of SD and the two days following the 2 nights of recovery sleep. Daily averages were statistically significant for reaction times on the second day of SD and for major lapses on both SD days and the day after the first night of recovery sleep. Therefore, the beneficial effects of this stimulation paradigm on performance extended for at least three days, with the STIM group back to baseline performance after only on night of recovery sleep. The SHAM group was still experiencing the deleterious effect of the restricted sleep/SD after the second night of recovery sleep. This demonstrates that the enhancement of slow oscillatory power with SO-tDCS, the presumed physiological effect of SO-tDCS, will effectively accelerate the restorative properties of a period of sleep, rendering a short period of sleep, in this case a two hour period, equivalent to a substantially longer period of natural sleep in terms of its ability to dissipate homeostatic sleep drive and return cognitive performance closer to a baseline level of performance that would be experienced with a full night of sleep that is effective at completely eliminating the sleep drive accumulated prior to that night of sleep. We utilized a very short sleep restriction period in order to accentuate the difference in sleep dissipating effects of natural sleep + SO-tDCS versus natural sleep alone (see further discussion of this issue below), and stimulation only occurred during 50 minutes of that two hour sleep period. We suspect that a longer period of stimulation during a two hour sleep period would have further accelerated the sleep drive dissipation process, producing even greater resilience to the deleterious evvectsof sustained waking on performance. What would be the shortest period of sleep enhanced by SO-tDCS that could completely eliminate homeostatic sleep drive would have to be assessed in future studies.

Insufficient sleep represents a major public health issue facing today’s society, in which access to vast repertoire of information and entertainment is available 24 hours a day and increasing constantly, with an ever greater percentage of the population progressively sleeping less and less per night, leading to tremendous individual and societal costs in terms of performance and productivity, as well as mental and physical health consequences (Chattu et al, 2018). Given the hypothesized mechanistic roles that have been attributed to sleep slow waves in both sleep related memory consolidation and sleep restorative function in the past 15 to 20 years, it is not surprising that there has been significant enthusiasm directed toward slow wave enhancement during sleep. Due mainly to issues related to side effects and tolerance as well as the somewhat modest level of SW enhancement observed in pharmacological trials (Walsh et al, 2005; 2010; Baandrup et al, 2026), enthusiasm for pharmacological slow-wave enhancement has waned. Previous research using pharmacological agents or acoustic stimulation to enhance slow-wave activity and assess the enhancement sleep’s restorative properties have demonstrated modest benefits at best. The majority of slow-wave enhancement studies have utilized acoustic stimulation or transcranial electrical stimulation to assess the effect of slow wave enhancement on sleep related memory consolidation. With respect to oscillatory transcranial stimulation, the results have been somewhat mixed (Feher, 2021). Marshall and colleagues (2006) developed the slow oscillation tDCS stimulation paradigm, consisting of an oscillation at the slow oscillation peak frequency of purely positive current, unlike transcranial alternating current stimulation (tACS), which oscillates between cycle phases of positive and negative current. The theoretical motivation for this novel oscillation paradigm was presumably to combine the benefits of a pure DC stimulation which Marshall theorized would induce a negative DC shift in the scalp electrical potential, which itself leads to an enhancement of SO power (Marshall, 2004) and the entraining properties of the superimposed SO frequency oscillation to further enhance SO Power. This induced negative DC shift would enhance a physiological negative shift that occurs naturally in NREM sleep and fosters the induction of sleep specific oscillations, including sleep spindles and the slow oscillation (Marshall et al, 1996; 1998) However, because the initial beneficial effects on memory were modest (Marshall, 2006) and attempts to replicate the findings in healthy participants as well as younger and older participants as well as patient population yielded very mixed results, the efficacy of SO-tDCS with the very weak currents utilized by Marshall, as well as various groups attempting to demonstrate SO-TDCS efficacy for memory consolidation has been called into question. Some studies have replicated the modest memory enhancing effects found by Marshall and colleagues, while others have not (Feher, 2021)refs). One study even failed to find significant cortical SO entrainment with stimulation applied directly to the cortex of human participants (Lafon et al, 2017). Further work has suggested a dose response relationship of current magnitude to the magnitude of low frequency stimulation induced oscillatory enhancement, with even weak currents producing some entrainment of cortical neurons to the slow oscillation (Johnson et al, 2021), further adding to the debate. Our study is to our knowledge the first study to use SO-tDCS for the express purpose of evaluating this stimulation paradigm as a methodology to enhance the restorative properties of the SO and thus to enhance next day performance as opposed to assess its use to enhance sleep’s memory consolidation properties. The very robust and long lasting performance enhancement findings demonstrated in our study should put to rest the issue of the ability of weak oscillatory currents to induce sufficient SO activity enhancement to significantly accelerate the restorative properties of sleep. Critics of the beneficial effects of SO transcranial electrical stimulation with weak currents have argued that there is insufficient current to produce SO enhancement, though this remains a matter of debate. They argue for alternative explanations for the benefit of SO-tDCS on memory performance assessed immediately after sleep enhanced with SO-tDCS, which include a placebo effect and indirect effects of cranial nerve stimulation, but not an enhancement of endogenous slow oscillatory activity. The inability to document an increase in SO power due to stimulation based electrical artifact certainly adds weight to the skepticism about the proposed physiological basis of SO-tDCS administration. However, with respect to our findings, we do not regard it as plausible that any other mechanism than an increased number of SO cycles could be responsible for the sustained beneficial effect of fifty minutes of stimulation on the sustained resilience of waking performance, indicative of an enhanced reduction of homeostatic sleep drive during the 2 hour sleep period in the STIM versus the SHAM participants. That beneficial effect spanned all 46 hours of subsequent total sleep deprivation experienced by participants in our study and in fact persisted into the post-recovery sleep period. Notably, the relative beneficial effect on performance for STIM versus SHAM participants increased over the sleep deprivation period and were most pronounced during nadirs in the circadian drive for arousal, as would be expected by a true enhancement of sleep drive dissipation but would be difficult to attribute to any other mechanism, including placebo effect. SO activity, to our knowledge, is the only neurophysiological mechanism by which homeostatic sleep drive can be dissipated.

The slow oscillation stimulation waveform, as innovated by Marsahall and colleauges, designed to mimic the physiology of the endogenous slow oscillation, consists of 4 components: the UP and DOWN states as which are plateau and nadir states characterized by an average membrane potential of neurons in the thalamocortical system, about 20 mV apart. The other two components are the Up to DOWN and DOWN to UP state transitions. One cycle of the SO consists of one iteration of each of these four components. Each of these components individually has been hypothesized to underlie one or more mechanistic bases for slow oscillation’s role in sleep’s restorative processes, based on different hypotheses about the underlying physiological basis for those properties. It is currently unclear which component or combination of components is involved and to what degree in the restorative properties that the SO contributes to sleep. None of the mechanistic hypotheses or the way in which a specific component of the slow oscillation may contribute to them is in any way mutually exclusive of the others.

Already mentioned as a prevalent theory of the restorative property of sleep is the synaptic homeostasis hypothesis, or SHY, which postulates that sustained waking is associated with ever increasing global synaptic strength of the cerebral cortex and that sleep and the SO in particular actively downscales synaptic strength to return the cortex to a more efficient state, with more salient increases in synaptic strength somehow preserved relative to less salient ones. Tononi and Cirelli (2006) proposed that the high firing rate of cortical neurons in the UP state, and in particular at the end of the UP state, followed by the abrupt transition to the DOWN state with a complete absence of firing would lead to a reduction in the strength of synapses of neurons connected to the neurons firing prior to the end of the UP state since an absence of firing of post-synaptic neurons in the brief period after firing of a presynaptic neuron leads to long-term depression of that synapse via the mechanism of long-term depression. In this case, it is the abrupt, synchronized UP to DOWN state transition that underlies the mechanistic basis of synaptic depression that could contribute to an overall sleep-related reduction in global synaptic strength.

A series of recent studies have also contributed evidence that the UP state of the SO may provide a mechanistic substrate to downscaling of synapses (Gulati et al, 2017; Bartram et al, 2017; Norimoto et al, 2018; Gonzalez-Rueda et al, 2018). Of note, one difference between the characteristics of neuronal activity during the UP state of the sleep SO and during wakefulness is there is a much greater occurrence of burst firing in the UP state versus during wakefulness (Ohyama et al, 2020). As bursts from a pre-synaptic neuron will tend to dramatically increase the likelihood of firing of post-synaptic neurons, burst firing in a specific subset of appropriately tagged neurons will help ensure the maintenance of synaptic strength in salient synapses, while the majority of synapses are undergoing a process of down selection.

Levenstein et al (2017) have provided a variation on SHY that implicates the DOWN to UP state transition in sleep’s restorative function. According to this theory, sustained wakefulness results in an ever-increasing disparity between the firing rates of high firing rate neuronal and low firing rate neurons. That is, neurons with higher firing rates increase their rates due to increased synaptic strength and the opposite effect is observed in lower firing rate neurons. Sleep and the SO leads to a very specific type of synaptic downscaling based on the order of neuronal firing during the DOWN to UP state transition.

Finally, a few hypotheses about the physiological basis of sleep’s restorative properties attribute mechanistic importance to the DOWN state specifically. One such proposal is that sleep is necessary for cellular maintenance and repair processes to take place to prevent irreversible damage to neurons that might ultimately result in cellular apoptosis (Scharf, 2011). Such repair optimally requires an absence of neuronal firing. Given the massive connectivity among cortical neurons, to attain a state of neuronal silence requires synchronized neuronal silence among all or nearly all neurons, a state that exists in the SO DOWN state (Vyazovskiy and Harris, 2013). DOWN states typically have a duration of a few hundred milliseconds. More prolonged DOWN states might be more efficient in completing the repair function of sleep, but presumably the alternation between UP and DOWN states balances the information processing functions of NREM sleep that may require ample neuronal firing in the UP state with the restorative repair function that depends on the DOWN state. Interestingly, the recovery sleep of sleep deprived animals is characterized by a lengthening of the average DOWN state duration by about 25 percent (Hajnik et al, 2013), as well as an increase in the overall frequency of DOWN states, at the expense of UP state durations, which are shortened, an indication of the importance of DOWN states to the sleep restorative process.

To summarize the current knowledge of the physiology of sleep restorative processes: there appears to be a number of mechanisms by which all four of the components of the SO cycle individually contribute to the dissipation of sleep drive, but to date no other viable physiological mechanisms have been proposed. The importance of transitions to and from SO DOWN states is so essential to brain function and likely neuronal survival, that such activity “leaks” into wakefulness when its magnitude is insufficient due to limited sleep. In fact, waking EEG activity under conditions of sufficient sleep pressure can become almost indistinguishable from true NREM sleep due to the abundance of intrusive DOWN states (Vyazovskiy et al, 2009). We regard this as strong evidence that SO-tDCS did indeed increase SO activity in our stimulated participants in order account for the robust resilience these participants exhibited to the detrimental effects of sleep deprivation on performance, only attributable to an accelerated dissipation of sleep drive during the 2 hour period of sleep.

In this “proof of concept” study, we attempted to maximize the likelihood of detecting a beneficial effect of stimulation on subsequent performance by utilizing a very short period of sleep opportunity (2 hours), to ensure the restorative effects of natural sleep in that period were insufficient to satisfy the homeostatic sleep drive that had accumulated over a normal 16 hour period of daytime wakefulness, so that any beneficial effect of stimulation on the restorative process would be accentuated compared to the sham participants who would only have experienced a fraction of complete sleep drive elimination. Such benefit would also be accentuated by the prolonged period of sustained wakefulness after the restricted sleep opportunity, with to the residual sleep drive remaining after the restricted sleep period in combination with the ever-accruing homeostatic sleep drive of 46 continued hours of wakefulness, which would be particularly problematic during the nadir of the circadian rhythm in the early morning in the absence of the circadian drive for arousal. We would argue, for example, that performance after stimulation during part or even all of a full eight hours of nocturnal sleep compared to sham stimulation or baseline performance would likely demonstrate little or no improvement as the endogenous SO in a healthy sleeper would likely have satisfied essentially all of the accumulated homeostatic drive for sleep restoration in such an individual without a pre-existing sleep debt (which we attempted to eliminate with our study design). We additionally theorized that a beneficial effect of SO enhancement during recovery sleep after a prolonged period of sleep deprivation would likely be minimal as endogenous SO activity is enhanced naturally after sleep deprivation by as much as 100 percent and therefore SO-tDCS enhanced SO power would likely not significantly exceed the SO power during natural recovery sleep (Lamond et al, 2007). Therefore, we suspect that SO enhancement with SO-tDCS will be most beneficial or perhaps only beneficial during restricted periods of sleep as we demonstrated. Of course, individuals typically experience extended or chronic periods of recurrent sleep restriction over many days, weeks, months or longer. Would SO enhancement with stimulation be beneficial over multiple consecutive nights of restricted sleep? One might argue that, as in recovery sleep after total sleep deprivation, a natural compensatory increase in SO power during subsequent nights of restricted sleep opportunity would occur given the increasing accumulation of homeostatic sleep drive that is not being adequately dissipated due to insufficient sleep time during the restricted periods of sleep. Such a significant endogenous increase in SO power would likely significantly limit the value of exogenous enhancement of SO activity and potentially limit the benefits of SO enhancement to a single period of sleep restriction in the absence of chronic sleep debt, likely a less common real life sleep issue. However, such a dramatic increase in SO power as is seen in recovery sleep after total sleep deprivation (TSD( does not appear to occur in the setting of chronic sleep restriction, despite a sustained and significant waking performance decrement. Contrary to the increases in SO power that occurs during recovery sleep after TSD, SO power during multiple consecutive nights of sleep restriction only increases by only about 0 to 20 percent (Skorucak et al, 2018; Voderholzer et al, 2011; Van Dongen et al, 2003), presumably leaving sufficient opportunity for additional exogenous SO enhancement.

However, why is it that sleep SO power during chronic sleep restriction is only minimally increased when that level of NREM SO power remains insufficient to fully dissipate homeostatic sleep drive, as demonstrated by the negative impact of chronic SR on performance? One likely reason appears to be that a major component of compensatory increase in slow-wave activity (DOWN states) occurs during wakefulness rather than during the restricted sleep periods, as alluded to previously (Leemberg et al, 2010). We believe that in the short term (recovery sleep after a single period of TSD), maximization of sleep restorative processes is prioritized over maintenance of normal sleep micro and microarchitecture, resulting in the pronounced increase in recovery sleep SO power. However, on a more chronic basis with nightly or near nightly sleep restriction, we believe that such sleep architecture maintenance is prioritized. As a result, and slow-wave DOWN states begin to intrude into wakefulness, necessitated by the need for restorative processes in the absence of sufficient sleep, compensating for the lack of sleep SO. It has been definitively demonstrated that SW activity during wakefulness (isolated DOWN states in the theta or delta EEG band frequency range) can compensate for sleep SO activity and indeed reduce the drive for SO activity during subsequent sleep. That is, SO/DOWN state activity during wakefulness serves a restorative function (Driessen et al, 2026; Vatikutti et al, 2026). Optogenetic activation of somatostatin-positive interneurons resulting in the occurrence of isolated DOWN states during wakefulness in a specific region of cortex significantly decreases SO power during subsequent sleep (Driessen et al, 2026). However, this “leakage” DOWN state activity into wakefulness, which primarily occurs locally rather than globally in the cortex, while successfully reducing homeostatic sleep drive, is associated with a sustained interruption in neuronal firing for up to several hundred milliseconds, resulting in the significant a performance decrement that oyccurs during chronic sleep restriction. Exogenous enhancement of sleep SO power with SO-tDCS during chronic sleep restriction would theoretically reduce or eliminate the drive for DOWN state activity during wakefulness and improve or potentially normalize waking performance. For that reason, we foresee SO-tDCS as having a role in the maintenance of waking performance in the setting of persistent sleep restriction (over multiple nights). Future studies consisting of multiple consecutive nights of restricted sleep supplemented with SO-tDCS will be able to ascertain the efficacy of such a therapy.

We have previously noted that the efficacy of SO-tDCS to induce SO activity and enhance SO based functions has been called into question since the publication of the seminal study by Marshal et al (2006) demonstrating an SO-tDCS based enhancement of sleep-related memory consolidation due to a lack of consistent replication of this effect in subsequent studies, with a few studies successfully demonstrating the modest memory effect in healthy participants as well as patient populations but with a number of studies unable to replicate this effect in healthy participants of various ages (Feher et al, 2021).

The fundamental mechanism for SO-tDCS based memory enhancement is proposed to be entrainment of a significant component of cortical neurons to the exogenous rhythmic oscillation of applied current, essentially increasing the number of SO cycles, perhaps even eliciting a continuous oscillation at the peak frequency of the endogenous SO, 0.75 Hz. Exactly what percentage of cortical neurons oscillate with the endogenous SO is unclear. The original work by Steriade suggested the vast majority (greater than 90%) of both excitatory and inhibitory neurons in a region under study oscillated in tandem. More recent evidence suggests a much lower percentage of neurons in a region may oscillate with the SO (Niethard et al, 2018). With respect to memory consolidation, the SO is believed to enhance sleep related memory consolidation via the following mechanism (Steriade and McCarley, 2005). At the time of the cortical transition from the electrically silent DOWN state to the depolarized UP state, corticothalamic projection neurons initiate and synchronize the generator of spindle activity, the thalamic reticular nucleus (nRT), to fire bursts of action potentials at spindle frequency, which in turn elicits spindle bursts in thalamocortical projection neurons throughout the rest of the thalamus. The thalamus transfers this spindle frequency activity to apical dendrites of cortical neurons, which generate the scalp recorded EEG signal. The hyperpolarized DOWN state preceding the synchronized transition to the UP state ensures that all nRT neurons are silent, hyperpolarized and poised for synchronous bursting, once activated.

The SO, during its rapid transition from DOWN to UP state, simultaneously triggers spindles (thalamus) and sharp-wave ripple potentials (SWR, hippocampus) to transfer hippocampally encoded memories to neocortex in a distributed manner, a process known as “systems consolidation” (Lutz et al, 2015). High levels of calcium currents entering cortical pyramidal cell apical dendrites during spindles are essential to the induction of synaptic plasticity and to this memory consolidation process (Seibt et al, 2017). After days, weeks months or longer of recurrent reactivation of memories and transfer to cortex, hippocampally encoded memories may be completely transferred to neocortex, no longer requiring hippocampal links to cortex for memory activation and recall, per systems consolidation theory.

Presumably, an SO-tDCS induced continuous SO will substantially increase spindle density over endogenous levels, since there will be an increase in SO UP states, each of which is potentially associated with spindle generation, and may also optimize the phase relationship of spindles to SO: at the DOWN – UP state transition and early in the subsequent UP state, a phase relationship that optimizes the memory consolidation process (Ladenbauer et al, 2021). However, due to the electrical artifact induced by the stimulation, analysis of the EEG response to stimulation is problematic, especially at the SO stimulation frequency (typically 0.75 Hz), so that the effective enhancement of SO power must be inferred either from a behavioral response attributed to SO enhancement (for example, improved memory performance after sleep with versus without stimulation) or from indirect evidence of an increase in SO power during brief interstimulus intervals felt to represent the persistence of a continuous SO induced during the stimulation period. Critics of the efficacy of SO-tDCS regard these inferences as incorrect or poorly reflective of actual SO power during SO-tDCS (Lafon et al, 2017). In support of this latter viewpoint, a recent study (Lafon et al, 2017) attempted to assess the efficacy of SO-tDCS administered by scalp electrodes by recording cortical physiology directly with electrocorticography in patients undergoing invasive monitoring for epilepsy. This study assessedthe efficacy of the SO stimulation to enhance spindle power (presumably localized to the SO UP state), as the power of spindle frequency range EEG activity could be assessed, though an SO activity power assessment was not possible due to electrical artifact. Spindle frequency power served as a surrogate for SO power as well as representing a physiological process necessary for systems consolidation to occur. Despite successfully recording spindle activity during SO UP states during NREM sleep at baseline, during SO-tDCS the recordings were essentially devoid of spindle frequency activity, although the presence ofUP states could not be specifically identified. This finding suggested an ineffective exogenous induction of SO activity which was attributed to an insufficient magnitude of electrical current generated by the very weak fields utilized in the study. The investigators concluded that SO-tDCS at standard intensities administered in published studies was insufficient to induce SO activity in the cerebral cortex. Any beneficial effects of the stimulation procedure on memory performance would have to be explained by another mechanism, including potentially a placebo effect, the investigators concluded.

However, several points about the relationship of SO mediated memory consolidation and SO mediated sleep restorative processes are important to consider in light of the purpose of our study, to specifically enhance sleep restorative properties, and its results. 1) spindle oscillations are not ubiquitous during SO UP states. The restorative function attributed to the SO is largely independent of and may be dissociated from sleep related memory consolidation. 2) We propose that SO-tDCS may induce SO activity that is devoid of spindle activity 3) Sleep-related memory consolidation processes mediated by SO physiology occur independent of spindle physiology.

Point 1: Firstly, General anesthetics produce sleep-like slow waves that do not support normal spindle physiology during the depolarized phase of the waves, since these anesthetics induce a level of thalamocortical membrane potential hyperpolarization that does not support such spindle physiology, and yet anesthesia induced slow waves exhibit sleep-like restorative processes (Tung et al, 2004) Light anesthesia induce by propofol does produce alpha frequency oscillations similar to spindles, but with a unique physiology that does not support the synaptic plasticity properties of spindles (Vijayan et al, 2013). Secondly, Bernardi et al (2018) categorize slow waves into two types: type I, consisting of global high amplitude DOWN states unassociated with UP state spindle generation, and Type II, consisting of lower amplitude DOWN states that trigger UP state spindles. Both types are believed to contribute to sleep restorative processes. Thirdly, and finally, Ngheim et al (2020) describe a slow wave physiology continuum based on the underlying level of cortical cholinergic tone, which determines the polarization level of the thalamocortical system. At one end of the continuum exists a rhythmic SO with short UP/DOWN states of similar duration, without UP state spindle generation, associated with lower cholinergic tone and deeper NREM sleep. The other end of the continuum consists of SO activity with longer and more variable duration UP than DOWN states, during lighter NREM sleep, frequently associated with UP state spindle activity. Notably, the higher frequency, rhythmic SO activity without spindles is regarded as possessing greater sleep homeostatic drive dissipating restorative function than the more irregular SO cycles found in lighter NREM associated with spindle activity. Clearly, one cannot conclude that the absence of spindle physiology during a period of NREM sleep is definitive evidence of the absence of SO physiology with restorative properties. To the contrary, the absence of spindles may actually be beneficial for the sleep restorative function of the SO, and it has been suggested that elimination or suppression of spindles may facilitate that restorative function.

Point 2. We propose that tDCS, or in this case the 260 µamp current plateau of the SO-tDCS, essentially a brief period of sustained tDCS, may actually lead to the suppression of sleep spindles during the UP state of the SO induced by the stimulation. The current scientific consensus on the role of tDCS on polarization of cortical neurons is that the effect is neuronalc ompartment dependent, e.g. axon versus soma versus apical dendrite (Rahman et al, 2013; 2015; Jackson et al, 2016; Aberra et al, 2023). Layer 5 (L5) pyramidal neurons exhibit the greatest degree of tDCS induced compartment polarization due to their profound length. Specifically, they possess long apical dendrites such that the neuron spans most layers of the cortex and generates an electrical dipole of large magnitude: with the cell soma located in layer 5, basal dendrites in layers 5 and 6 and with apical dendrites projecting up to supragranular layers 1 – 3, with a tuft of superficial dendrites located in these layer. Importantly, tDCS is believed to depolarize the soma and basal dendrites and hyperpolarize the majority of the apical dendrites and dendritic tuft (Rahman et al, 2013; 2015). This represents an active “sink” of positive current entering the soma in L5 and an active “source” of positive current existing the apical dendrites in the supragranular layers. These layer 5 pyramidal neurons appear to orchestrate the slow oscillation with its transition between UP and DOWN states (Neske, 2016; Chauvette et al, 2010) and their apical dendrites primarily generate the electrical oscillations of cortical spindle activity giving rise to EEG spindles (Seibt et al, 2017). To the degree that an oscillation induced by SO-tDCS oscillates in relative synchrony with the exogenous applied current, the current plateau segment of the SO-tDCS would correspond to the UP state, and this depolarized state with increased firing of L5 neurons would be consistent with an induced SO UP state. This applied current would help support the bursts of action potentials fired by L5 pyramidal neurons during the endogenous UP state (Ohyama et al, 2020). However, it is clear that a state of hyperpolarization of the apical dendrites in supragranular layers induced by the anodal current would be inconsistent with dendritic physiology necessary to support the emergence of sleep spindles during the UP state, during which L5 pyramidal neuronal apical dendrites are depolarized compared to waking levels due to intense inhibition of the somatostatin positive (SOM+) interneurons populating layers 1 – 3 that exert inhibitory tone over these apical dendrites. The silencing of these SOM+ neurons during the UP state fosters the elicitation of spindle physiology (Seibt et al, 2017). We suggest that the polarization state of these neurons induced by tDCS is strikingly similar to that found during the “solitary slow oscillation,” without UP state spindles, as depicted in Fig. 6 of Niethard et al (2018). Under this natural endogenous condition of SO cycles without spindles, apical dendrites are hyperpolarized due to intense activity in SOM+ inhibitory interneurons located in supragranular layers while the soma is depolarized due to reduced inhibition by parvalbumin positive (PV+) interneurons in the infragranular layers, which regulate L5 pyramidal neuron somatic inhibitory tone. We believe this strong dipolar electrical neuronal state (negative end of the dipole in supragranular layers /positive end in infragranular layers) would foster NREM restorative processes by limiting UP state duration and increasing SO frequency within the natural SO range, similar to SO activity during deep sleep (Nghiem et al, 2020) or recovery sleep after sleep deprivation, which is characterized by an increase in SO power, a decrease in SO frequency (due to a decrease in UP state duration) and a decrease in SO associated UP state spindle activity (Olbrich et al, 2014; Bersagliere et al, 2010). Such SO activity, with an absence of spindles, while clearly not supporting the spindle related systems consolidation process, likely enhances sleep restorative processes. The “slow oscillation + spindle state” in Niethard Fig. 6 depicts disinhibited (depolarized) apical dendrites due to strong SOM+ interneuron inhibition and a reduced level of somatic depolarization due to relative PV+ interneuron activation. This “SO + spindle state” would support a SO UP state current source density (CSD) profile similar to the one recorded in human epilepsy patients undergoing invasive monitoring with cortical multi-electrode depth recordings during a period of spindle rich SO activity: consisting of a very strong current “sink” in supragranular layers associated with strong calcium currents depolarizing apical dendrites of L5 pyramidal neurons, along with a weak sink in infragranular layers due to the weaker currents depolarizing the soma, under a state of relative inhibition (Csercsa et al, 2010).

Therefore, we propose that the SO induced by SO-tDCS may indeed be devoid of sleep spindles in the setting of robust transition between UP and DOWN states mediating sleep restorative processes which would be consistent with the finding of Lafon et al of an absence of sleep spindles in human recordings with electrocorticography during SO-tDCS. In fact, we believe that the presence, absence or magnitude of various SO physiological features likely represent a trade-off between values that optimize memory consolidation processes versus restorative processes, which may be prioritized by brain physiology differently based on past sleep history, depth of NREM sleep, pre-sleep waking behavior (exposure to salient information to be remembered), and other factors. Based on this conceptualization, we suggest that SO-tDCS, with the specific stimulation parameters utilized in this study, will induce SO activity with physiology that may optimize SO based restorative function and may actually relatively minimize physiology to support sleep-related systems consolidation, namely UP state generated sleep spindle activity.

Point 3: In our study, we were able to demonstrate a beneficial effect of the SO-tDCS stimulation of memory performance after the restricted sleep period, essentially replicating the effect of Marshall et al (2006), but with a much shorter period of sleep (2 hours versus 7.5 hours). The discussion of point 3 would certainly raise the question of how we demonstrated a beneficial effect of 50 minutes of SO-tDCS on the memory for word pairs (PWAT) learned immediately prior to the sleep period compared to sham stimulation. If our SO-tDCS paradigm was actively suppressing UP state spindles, the sham participants experiencing natural sleep would likely have produced a substantially greater spindle density during that second hour of sleep. We believe that this finding is due to the presence of an SO induced enhancement of a focal memory consolidation process of engrams encoded within the hippocampus – that is, a strengthening of the hippocampally encoded memory, rather than due to a systems consolidation or transfer of memory to a distributed neocortical representation. To support successful systems consolidation, in the very short term (minutes to a few hours), the hippocampally encoded component of all declarative memory must be consolidated so that a transfer of information can then take place over days to weeks to months. A number of studies have demonstrated how the cortical SO, in addition to stimulating hippocampal ripple activity to reactivate memory information in cortex, also activates local sharp wave-ripple activity within the hippocampus to stabilize and organize newly encoded memories, strengthening synapses within CA3, and CA3 – CA1 connections, that are part of a memory engram (Rolotti et al, 2022; Sadowski et al, 2016; Van den Ven et al, 2016). Additionally, the SO activates “dentate spikes,” brief potentials comprised of dentate gyrus granule cell activity that reorganizes memory engrams, including the merging of neural assemblies of highly related memories (Nokia and Penttonen, 2022). These are memory consolidation properties intrinsic to the hippocampus that are fostered by SO activity that does not involve spindle physiology. Regardless, our findings confirm the concept that SO stimulation enhances sleep related consolidation of episodic memories.

There is some debate over the nature of episodic memories that are permanently transferred from the hippocampus to the cortex (Moscovitch et al, 2016). One hypothesis is that all declarative memories including all episodic memories, regardless of the level of detail, are eventually completely transferred to cortex with no residual activity in the hippocampus. This seems to be the version of the theory favored by a number of investigators using SO-tDCS for the purpose of enhancing sleep-related memory consolidation. An alternative theory is the that predominantly generalizable information is transferred to cortex (Sun et al, 2023), consistent with the notion of “interleaved learning” of semantic memories (McClelland et al, 1995). Common information is extracted from multiple episodic memories that are reactivated recurrently over an extended period of time (weeks, months or longer) and such extracted information is stored in cortex, in the form of semantic memory or schemas. In this conceptualization, highly episodic knowledge containing detailed episodic information would maintain a representation in the hippocampus, which would permanently be involved in the co-activation of the memory specific information that is distributed widely in cortex. Only the basic aspects of a detailed episodic memory, known as a gist memory, may be ultimately stored primarily in cortex (ventromedial prefrontal cortex) but even gist memories are not be conceived to be completely devoid of a hippocampal link (Robin et al, 2017; Sekeres et al, 2018). For episodic memory rich in details, it is unclear what if any consolidative processes would take place in cortex subsequent to encoding and consolidation within in the hippocampus. Rather, for these memories, a robust engram would be maintained within the hippocampus to ensure the ability to co-activate these rich details concurrently as part of a vivid episodic memory. It has been suggested, in fact, that there are mechanisms in place to prevent “unregulated neocortical memory transfer” from the hippocampus (Sun et al, 2023) that would be detrimental to knowledge generalizability as would be likely in in the case of the word associations learned during the declarative memory task in our study, the PWAT task. Such novel associations of semantically unrelated words could be regarded as “outlier information” that would not add to generalizable knowledge of the world. In such cases, local hippocampal consolidative processes would likely dominate over a “systems consolidation” based transfer of information to cortex, probably supplemented by consolidate changes involving direct hippocampal-cortical connections. In fact, we suggest that the memory of PWAT word associations are likely maintained in the hippocampus indefinitely (as long as they are remembered at all), as would be the case with all memories for which highly specific detailed aspects are recalled We regard the memories of the generally poorly associated lexical items comprising our PWAT task as highly episodic and therefore permanently dependent on hippocampal links rather than a purely cortical set of links among these semantic concepts. We regard it as far less likely that a significant contribution of hippocampal-neocortical transfer would transpire within a brief two hour period of post-learning sleep versus consolidation within the hippocampus of these newly encoded memories, especially those of such a highly episodic nature, which may in fact never truly undergo a process of hippocampal-neocortical transfer at all. Therefore, a potential absence of spindles in association with SO-tDCS, as suggested by the Lafon et al. study would have had little or no impact on stimulation induced improvement in highly episodic memory recall performance in our study, which we believe is primarily mediated locally in the hippocampus, with the exogenous SO significantly enhancing localized SW-Rs that “stabilize” the newly encoded episodic memories of the PWA word pairs. The limited selective activation of local SW-Rs and dentate spikes in the absence of spindles could conceivably be more beneficial to the local hippocampal consolidation process than what would occur in spindle rich NREM SO activity.

The following is a summary of our conceptualization of the potential for the use of SO-tDCS during periods of restricted sleep opportunity. SO-tDCS enhances sleep’s restorative properties and can dissipate accumulated homeostatic sleep drive significantly more efficiently than natural sleep and therefore promotes resilience to sustained wakefulness related performance decrements.

Though we only demonstrated such a benefit for stimulation during a single period of sleep restriction and a single subsequent period of wakefulness, we believe a beneficial effect is achievable over a series of consecutive nights due to the fact that the natural response of the brain to chronic sleep restriction is to maintain natural sleep architecture and transfer much of the demand for restorative slow wave activity to wakefulness. This waking slow wave significantly degrades waking performance. The exogenous enhancement of SO activity with SO-tDCS could satisfy a substantial component of sleep homeostatic drive and relieve the need for waking slow-wave activity, potentially indefinitely.

However, depending on the percentage of NREM sleep duration of the restricted sleep period during which an individual receives SO-tDCS, the stimulation paradigm will result in a significant alteration in sleep micro– and macro-architecture. For example, there may be a significant alteration in spindle density and the phase relationship of spindles to the slow oscillation, if indeed SO stimulation has a suppressive effect on UP state spindle activity. There is presumably a teleological reason that the brain prioritizes maintenance of sleep architecture at the expense of waking performance during chronic sleep restriction. Spindles not only play a crucial role in memory systems consolidation, but it is suggested that spindles arising in isolation in the absence of SO physiology are involved in the reorganization of already established cortical representations of information, likely crucial to the maintenance of an accurate and updated world model for predictive processing and survival (Friston, 2010). Additionally, recent evidence suggests sleep spindles have an important role in the adaptation to stress, likely an important function during a period of life characterized by chronic sleep deprivation (Natraj et al, 2023a; 2023b; Kleim et al, 2016).

Based on the above discussion, including the fact that long term inhibition of spindles and other potential alterations in sleep micro-architecture may potentially be induced by SO-tDCS, we foresee the potential of recommending the use of SO-tDCS during a substantial percentage of NREM sleep of a restricted sleep opportunity for a continuous period of several days or perhaps a few weeks, not for months or longer, should its efficacy over multiple nights be verified.

Certain limitations to our study should be emphasized. First, the study had a small sample size with a total of 27 participants. This was primarily due to the significant resource demands of a single study session, which included a total of 7 nights in the sleep lab; for the last consecutive 5 days/nights of the study, participants were confined to the lab for the full 24 hours. Optimally, the findings of this study will be replicated with a larger sample size in the future. Additionally, we initially planned a crossover design to reduce inter-subject variability, with all participants receiving SO stimulation and sham stimulation in a randomized order. However, due to the time demands of the study as just described, we found that recruiting participants who would be able to commit to two demanding study sessions within a reasonable period of time would be prohibitive and we therefore opted for a parallel study design.

Finally, we have already commented on the issue of the inability to quantitatively assess the effect of the SO-tDCS stimulation on SO power due to the electrical artifact that saturates the EEG amplifiers. Our conclusion that the stimulation was successful at enhancing SO power was necessarily inferred from the behavioral data demonstrating a robust beneficial effect of SO-tDCS on subsequent waking cognitive performance. For similar reasons, our hypothesis regarding the potential effect of SO-tDCS on sleep spindle activity remains speculative, though we believe it is supported by the underlying neurophysiology.

## Acknowledgements

This research benefitted greatly from discussions between JDH and Lisa Marshall regarding the SO-tDCS methodology and with Eric Wassermann and Maron Bikson regarding general transcranial electrical stimulation methodological issues. The authors would like to thank Samantha M. Riedy for her contribution to the analysis of our PWAT data.

## Disclaimers

The views expressed in this paper reflect the results of research conducted by the authors and do not necessarily reflect the official policy or position of the Defense Health Agency, the Department of War or the U.S. Government.

The identification of specific products or scientific instrumentation is considered an integral part of the scientific endeavor and does not constitute endorsement or implied endorsement on the part of the authors, DoD, or any component agency.

The study protocol was approved by the Walter Reed Army Institute of Research Institutional Review Board in compliance with all applicable Federal regulations governing the protection of human subjects.

This material has been reviewed by the Walter Reed Army Institute of Research. There is no objection to its publication.

The investigators have adhered to policies for protection of human subjects as prescribed in AR 70-25.

